# Cytosolic mannosyltransferases involved in the ER N-glycosylation pathway exhibit bacterial ancestry

**DOI:** 10.1101/2025.10.15.682550

**Authors:** Si Xu, Shuai Chen, Yu-He Gu, Yi-Fan Huang, Hideki Nakanishi, Xiao-Dong Gao

**Affiliations:** Key Laboratory of Carbohydrate Chemistry and Biotechnology, Ministry of Education, School of Biotechnology, Jiangnan University, Wuxi 214122, China; State Key Laboratory of Biopharmaceutical Preparation and Delivery, Institute of Process Engineering, Chinese Academy of Sciences, Beijing 100190, China

**Keywords:** activity assay, ALG glycosyltransferases, evolutionary trajectories, N-glycosylation pathway, phylogenetic analyses

## Abstract

N-glycosylation in eukaryotes begins with the assembly of a lipid-linked oligosaccharide on the endoplasmic reticulum membrane. As a pivotal post-translational protein modification, it is conserved across all three domains of life. However, the evolutionary origins of the N-glycosylation pathway remain a subject of ongoing debate in evolutionary biology, largely due to the limited availability of robust data regarding the evolutionary trajectories of the glycosyltransferases involved in this process. Here, we present phylogenetic analyses of the eukaryotic ALG1 and ALG2 mannosyltransferases (MTases), which are crucial for constructing the core trimannosyl Man3GlcNAc2 structure conserved in eukaryotic N-glycans. Our comprehensive phylogenetic study, combined with functional and structural analyses, suggests that the ALG2 MTase likely originated from a bacterial ancestor. This inference is further supported by the identification of sequential and functional ALG1 homologs exclusively within bacterial lineages, rather than in Asgard archaea or other archaeal groups. Our findings challenge the prevailing hypothesis that the eukaryotic N-glycosylation pathway primarily evolved from archaeal ancestors, instead suggesting a chimeric origin involving contributions from both bacterial and archaeal lineages.

**Synopsis:** 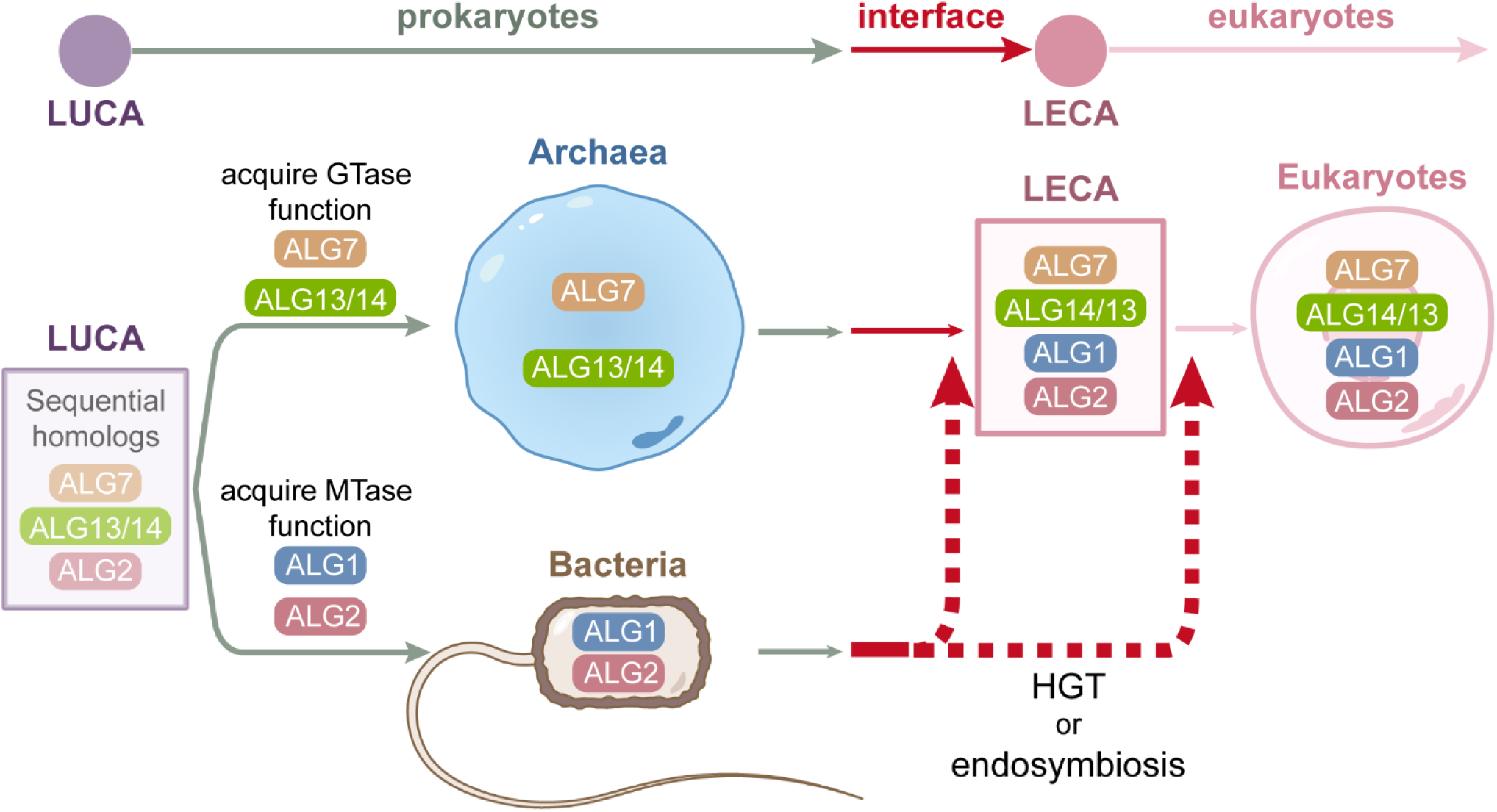

The ALG glycosyltransferase (GTases) involved in the initial stage of eukaryotic N-glycosylation on the cytosolic side of the ER membrane exhibit chimeric evolutionary origins. LUCA: Last Universal Common Ancestor; LECA: Last Eukaryotic Common Ancestor.

**Highlights:**

- The evolutionary trajectories of cytosolic ALG mannosyltransferases (MTases) were studied
- The ancestor of ALG2 exhibited in the LUCA but evolved its MTase activity in bacterial lineages
- The ALG1 MTase originated from bacterial lineages

## Introduction

In eukaryotes, N-glycosylation starts on the membrane of the endoplasmic reticulum (ER) with assembly of a precursor lipid-linked oligosaccharide (LLO) (Fig. 1A)(Aebi, 2013; Aebi *et al*, 2010; Schwarz & Aebi, 2011). Specifically, 14 sugars are sequentially attached to a polyisoprenyl lipid carrier (Fig. 1B) by membrane-associated ALG (Asparagine-Linked Glycosylation) glycosyltransferases (GTases)(Schmaltz *et al*, 2011). The LLO synthetic pathway exhibits distinct topological characteristics. Seven monosaccharides are sequentially assembled on the cytoplasmic face of the ER by two multi-GTase complexes (Fig. 1C). The initiation complex comprises the ALG7 UDP-GlcNAc phosphotransferase, in conjunction with the ALG13/14 heterodimeric UDP-GlcNAc transferase. These enzymes sequentially add GlcNAc-P and GlcNAc to the dolichol phosphate carrier (Dol-P), resulting in the formation of GlcNAc2-pyrophosphate-dolichol (GN2-PP-Dol) (Fig. 1C)(Lu *et al*, 2012; Noffz *et al*, 2009). Subsequently, the ALG1, ALG2, and ALG11 mannosyltransferases (MTases) function as a cytosolic MTase complex, transferring the subsequent five mannoses (M5) to the chitobiose core structure (GN2-) (Fig. 1C)(Gao *et al*, 2004; O’Reilly *et al*, 2006). The intermediate M5GN2-PP-Dol is then translocated across the ER membrane into the lumen by the RFT1 flippase(Chen *et al*, 2024; Helenius *et al*, 2002), where another seven sugars (four mannoses and three glucoses) are added to form the final G3M9GN2-PP-Dol precursor (Fig. 1A). Once assembled, the G3M9GN2 glycan is transferred en bloc to an asparagine residue within the Asn-X-(Ser/Thr) consensus sequence of target proteins by the oligosaccharyltransferase (OST) complex(Cherepanova *et al*, 2016; Mohorko *et al*, 2011; Shrimal & Gilmore, 2019). As the glycoprotein traverses the ER and Golgi apparatus, the protein-bound G3M9GN2 undergoes trimming and remodeling(Toustou *et al*, 2022). The resulting N-glycans exhibit greater structural diversity, yet all retain a core trimannosyl M3GN2 structure, which is the product of ALG1 and ALG2(Li *et al*, 2018; Xiang *et al*, 2022) (Fig. 1C). This characteristic establishes ALG1 and ALG2 as representative MTases for eukaryotic N-glycosylation.

**Figure 1.**
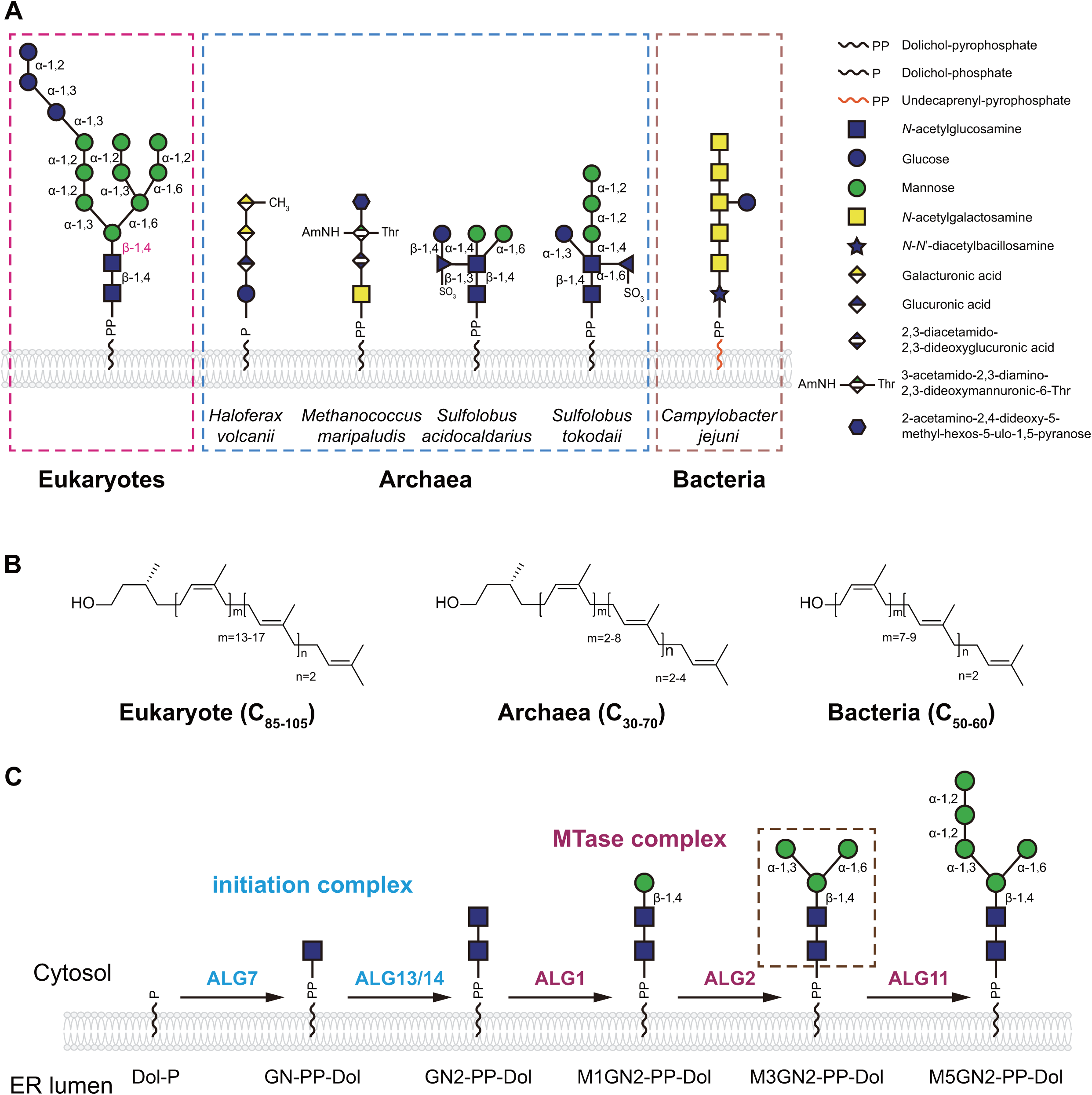
Comparison of lipid-linked oligosaccharides (LLOs) among the three domains of life, and the initial stage of the eukaryotic LLO synthesis at cytosolic surface. (A) The representative glycan structures of LLOs identified across the three domains of life. Eukaryotes possess a conserved tetradecasaccharide LLO. Archaeal LLOs exhibit considerable structural diversity, predominantly adopting linear configurations, although some species contain branched-chain structures. The LLO in *Campylobacter jejuni* (bacteria) features a linear glycan chain; (B) Diversity of the lipid carrier across the three domains of life. Eukaryotic organisms possess the longest fatty acid chains, ranging from 85 to 105 carbon atoms, whereas archaeal chains typically vary between 30 and 70 carbons, and bacterial chains generally contain 50 to 60 carbons; (**C**) Catalytic functions of cytosolic ALG glycosyltransferase (GTase) complexes involved in the early stages of LLO synthesis. The initiation complex consists of ALG7 and the ALG13/14 heterodimeric GTases. ALG1, ALG2, and ALG11 mannosyltransferases (MTases) constitute the MTase complex. The core trimannosyl M3GN2 structure is indicated by a dashed box.

N-glycosylation is a ubiquitous post-translational modification system found across all three domains of life(Nothaft & Szymanski, 2010; Weerapana & Imperiali, 2006; Yurist-Doutsch *et al*, 2008). In archaea, N-glycosylation occurs at the plasma membrane using dolichol-like lipids (Fig. 1B)(Yurist-Doutsch *et al*., 2008). The synthesized glycans are translocated from the periplasm to the outer leaflet of the plasma membrane, where the glycan moieties are primarily transferred to surface-layer proteins by OST homologs(Abu-Qarn *et al*, 2007). Archaeal N-glycosylation pathways generate a diverse array of LLOs across different species, which differ significantly from those found in eukaryotes (Fig. 1A)(Fujinami *et al*, 2017; Jarrell *et al*, 2014; van Wolferen *et al*, 2020). Bacteria also possess lipid-carrier-based N-glycosylation pathways. For instance, *Campylobacter jejuni* assembles a heptasaccharide onto undecaprenol pyrophosphate across the plasma membrane (Fig. 1A, B) and transfers it to specific asparagine residues of the target protein(Kawakami & Fujisaki, 2018; Knauer & Lehle, 1999; Wacker *et al*, 2002). Notably, prokaryotic N-glycosylation pathways involve the assembly of LLO intermediates across the plasma membrane, followed by their transfer to the target proteins via OST homologs — a process analogous to eukaryotic ER N-glycosylation(Abu-Qarn *et al*., 2007; Kowarik *et al*, 2006). These similarities suggest the existence of a potential common ancestral mechanism for the N-glycosylation pathway.

An early study suggested a closer phylogenetic relationship between archaeal and eukaryotic ALG7 homologs(Lombard, 2016). Consistent with this observation, several archaeal ALG7 homologs have been shown to functionally complement *alg7Δ* yeast cells, whereas bacterial homologs failed to do so(Meyer *et al*, 2017; Mikusova *et al*, 1996; Mitachi *et al*, 2016; Shams-Eldin *et al*, 2008). A recent study identified the Agl24 protein as the archaeal homolog of the eukaryotic ALG13/14 dimeric GTase(Meyer *et al*, 2022). Agl24 exhibits a closer phylogenetic relationship with the eukaryotic ALG13/14 GTase than its bacterial homolog MurG and, importantly, possesses the same β-1,4-GlcNAc transferase activity as ALG13/14 GTase. Given that ALG7 and ALG13/14 function as the initiation complex for ER N-glycosylation, these results contemplate an idea that the eukaryotic N-glycosylation pathway was inherited from archaea(Lombard, 2016; Meyer *et al*., 2022; Raval *et al*, 2022). Within the cytosolic MTase complex, ALG2 and ALG11 exhibit substantial sequence homology and are both classified within the GT4 family(Drula *et al*, 2022). Previous phylogenetic studies suggested a shared archaeal origin for ALG2 and ALG11(Lombard, 2016). Most recent updates on the phylogeny of the N-glycosylation pathway have identified sequential homologs of ALG1 in bacteria(Kifer *et al*, 2025). However, these findings require further validation.

The recent accumulation of genomic data has facilitated a more comprehensive understanding of the distribution of ALG GTases. In this study, we performed comprehensive analyses to elucidate the evolutionary history of human ALG1 and ALG2 MTases. A taxonomic homology survey revealed structural homologs of ALG2 across all three domains of life, suggesting its presence in the Last Universal Common Ancestor (LUCA). Further Bayesian clustering analysis, combined with functional investigations and structural modeling, provides strong evidence that the ALG2 MTase originated from a bacterial ancestor. Additionally, our analyses identified both sequential and functional ALG1 homologs exclusively within bacteria, indicating its bacterial origin. Taken together, our findings suggest that ER cytosolic ALG MTases exhibit bacterial ancestry, thereby challenging the prevailing hypothesis that the eukaryotic N-glycosylation pathway primarily originated from archaeal ancestors(Lombard, 2016; Meyer *et al*., 2022; Raval *et al*., 2022).

## Results

### BLAST analysis identifies cytosolic ALG MTase homologs across the three domains of life

To conduct a taxonomic homology survey, the *Homo sapiens* ALG1 (*Hs*ALG1) and ALG2 (*Hs*ALG2) were individually analyzed using the Basic Local Alignment Search Tool (BLAST) against the National Center for Biotechnology Information (NCBI) database (https://blast.ncbi.nlm.nih.gov/Blast.cgi/). BLASTp searches were carried out using an E-value cut-off of 1.00E-20 for eukaryotes and 1.00E-7 for prokaryotes. These relatively low E-values were selected to ensure a high level of confidence in identifying homologous relationships and especially, to differentiate between ALG2 and ALG11 homologs, which are considered as paralogs(O’Reilly *et al*., 2006).

A total of 4,654 homologous sequences were identified in eukaryotes for ALG2, comprising 187 from Protista, 2,312 from Fungi, 475 from Plantae, and 1,682 from Animalia ( Table 1; see details in Table EV1). Similarly, a total of 3,676 bacterial homologs were identified across all phyla described by the International Committee on Systematics of Prokaryotes (ICSP) from 2021 to 2023(Göker & Oren, 2023; Oren & Garrity, 2021; Oren *et al*, 2022). However, most prokaryotic sequences were primarily derived from the metagenome-assembled genome database, which limits our ability to ascertain the specific species from which they originate. In contrast to the widespread distribution of homologs in eukaryotes and bacteria, most archaeal homologs (a total of 862 sequences, see Table EV1) were identified within the TACK superphylum, with only a few sequences with higher E-values originating from the Euryarchaeota phylum, as well as the DPANN and Asgard superphyla. Compared to ALG2, a total of 4,839 ALG1 homologous sequences were identified in eukaryotes, comprising 208 from Protista, 2,364 from Fungi, 510 from Plantae, and 1,757 from Animalia (Table 1; see details in Table EV2). Furthermore, 204 bacterial sequences and only one archaeal sequence were identified as ALG1 homologs. Notably, this archaeal ALG1 homolog has a relatively high E-value of 5.00E-8, whereas the E-values for bacterial homologs range from 6.00E-22 to 1.00E-104 (Table EV2). This suggests that this archaeal sequence may not be a reliable homolog of ALG1 in this study and can be disregarded.

**Table 1.** Statistics of the number of ALG1 and ALG2 homologs.

| Protein | Total | Bacteria | Archaea | Eukaryotes (E value < 1.00E-20) |  |  |  |  |
| --- | --- | --- | --- | --- | --- | --- | --- | --- |
|  |  | (E value < 1.00E-7) | (E value < 1.00E-7) | Total | Protista | Fungi | Plantae | Animalia |
| ALG1 | 5044 | 204 | 1 | 4839 | 208 | 2364 | 510 | 1757 |
| ALG2 | 9192 | 3676 | 862 | 4654 | 187 | 2311 | 475 | 1681 |

From a comprehensive perspective, our BLASTp analysis of the cytosolic MTase complex provides the following evidence elucidating its evolutionary history: (1) the broad distribution of ALG2 homologs across all three domains of life suggests its presence in the LUCA; (2) the absence of reliable archaeal homologs implies a bacterial origin for the ALG1 MTase.

### The phylogenies of human ALG2 MTase reveal a closer evolutionary relationship with bacterial homologs

To further study the evolutionary pattern of ALG2 MTase, 63 proteins from above ALG2 homologs (Table 1, see Table EV3 for detailed sequences) were selected for multiple sequence alignment (MSA) followed by phylogenetic analysis. These sequence seeds comprise 16 from different archaeal lineages (including 9 from the top 25 hits of the TACK superphylum and 7 from the top 15 hits of the Asgard superphylum), 9 from different bacterial phyla or classes within the top 40 hits, 38 from eukaryotes (9 each from different model organisms of Protista, Plantae, and Animalia, and 11 from different model organisms of Fungi). The MSA results revealed that these 63 ALG2 homologs share a very high amino acids similarity (Table EV4). Noteworthy, the 7 sequences from the Asgard superphylum were intentionally included for phylogenetic study despite their relatively higher E-values compared to those of TACK homologs. The reason for selecting these sequences is that the Asgard superphylum has been considered as the closest prokaryotic relatives of eukaryotes(Liu *et al*, 2021; Spang *et al*, 2015; Zaremba-Niedzwiedzka *et al*, 2017).

To construct the Bayesian phylogenetic tree, a bacterial WbaZ sequence was used as an outgroup. This sequence corresponds to a capsular polysaccharide biosynthesis-associated protein belonging to the GT4 family, which also includes ALG2. As depicted in Figure 2, the initial sequences branching off from the outgroup are archaeal homologs. The remainder of the phylogenetic tree is partitioned into three distinct clades: one encompassing bacterial and protozoan clusters, another comprising plant and animal clusters (including several protozoan homologs), and an independent fungal clade. These findings suggest that bacterial homologs exhibit a closer phylogenetic relationship with eukaryotic ALG2. Notably, the archaeal branch can be internally subdivided into two groups: Asgard and TACK. Among these, TACK homologs sho w better similarity to bacterial and eukaryotic homologs (Fig. 2). In summary, the phylogenetic analysis reveals a close evolutionary relationship between eukaryotic ALG2 MTases and their bacterial counterparts. Our results provide evidence supporting the hypothesis that the ALG2 MTase originates from a bacterial ancestor.

**Figure 2.**
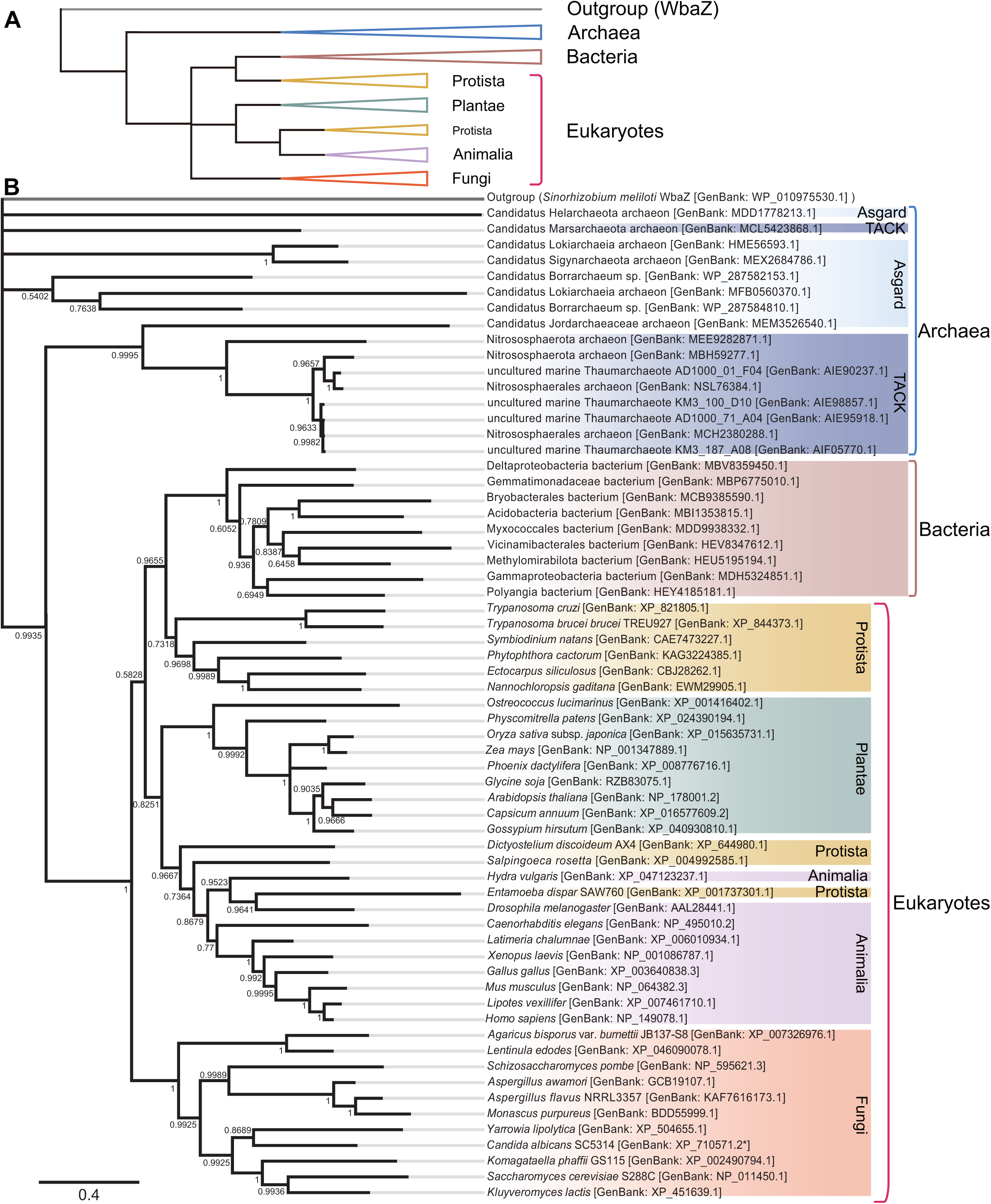
The bacterial ALG2 homologs exhibit a closer evolutionary relationship to those from eukaryotes than archaea. (**A**, **B**) Bayesian phylogenetic tree of ALG2 homologs was constructed from an alignment of 64 protein sequences generated using Clustal Omega. The background colors correspond to different kingdoms: Archaea (blue), Bacteria (brown), Protista (yellow), Fungi (orange), Plantae (green), and Animalia (purple). (**A**) provides a condensed overview of (**B**).

### Biological analyses identified functional ALG2 homologs in bacteria

The phylogenetic tree in Figure 2 is constructed based on DNA and protein sequence data, and it needs to be validated for reliability. To overcome the limitations of such phylogenetic approach, we tried to test the biological activity of sequential ALG2 homologs.

First, we conducted a yeast complementary assay using plasmid shuffle approach. *ALG2* homologous genes were cloned into a *2μ*/*LEU2* yeast expression vector and then transformed into a yeast *alg2Δ* strain, whose viability relies on the presence of a *URA3* plasmid-borne copy of *ALG2* gene from *Saccharomyces cerevisiae* (*ScALG2*) (see *Methods* for details). When cultured on media containing 5-fluoroorotic acid (5-FOA) to force loss of the *URA3-ScALG2* plasmid, only transformants with functional ALG2 homologs on the *LEU2* plasmid would allow *alg2Δ* growth (Fig. EV1A). For testing the complementation, 4 bacterial and 6 archaeal (2 from Asgard and 4 from TACK) ALG2 homologs were selected from the compilation of homologous sequences used for the above phylogenic analysis (Table EV3). After confirming the protein expression by western-blotting using anti-His tag antibody (Fig. EV1B), transformants were grown on the SD-Leu plate containing 5-FOA to remove the *URA3-ScALG2* plasmid. It was revealed that two bacterial homologs derived from Gammaproteobacteria [GenBank^#^: MDH5324851.1 (*Ga*ALG2)] and Polyangia species [GenBank#: HEY4185181.1 (*Po*ALG2)] respectively complemented the growth defect of *alg2Δ* cells, while the other two homologs derived from Deltaproteobacteria (*De*ALG2) and Methylomirabilota species (*Me*ALG2) failed to do so (Fig. 3A). In contrast to the bacteria, none of archaeal genes can complements the growth of *alg2Δ* strain, which include homologs derived from Thaumarchaeota KM3_187_A08 (*Th*ALG2-1), Thaumarchaeota AD1000_01_F04 (*Th*ALG2-2), Marsarchaeota (*Ma*ALG2), Nitrososphaerota (*Ni*ALG2), and Borrarchaeum (*Bo*ALG2-1 and *Bo*ALG2-2) (Fig. 3B).

**Figure 3.**
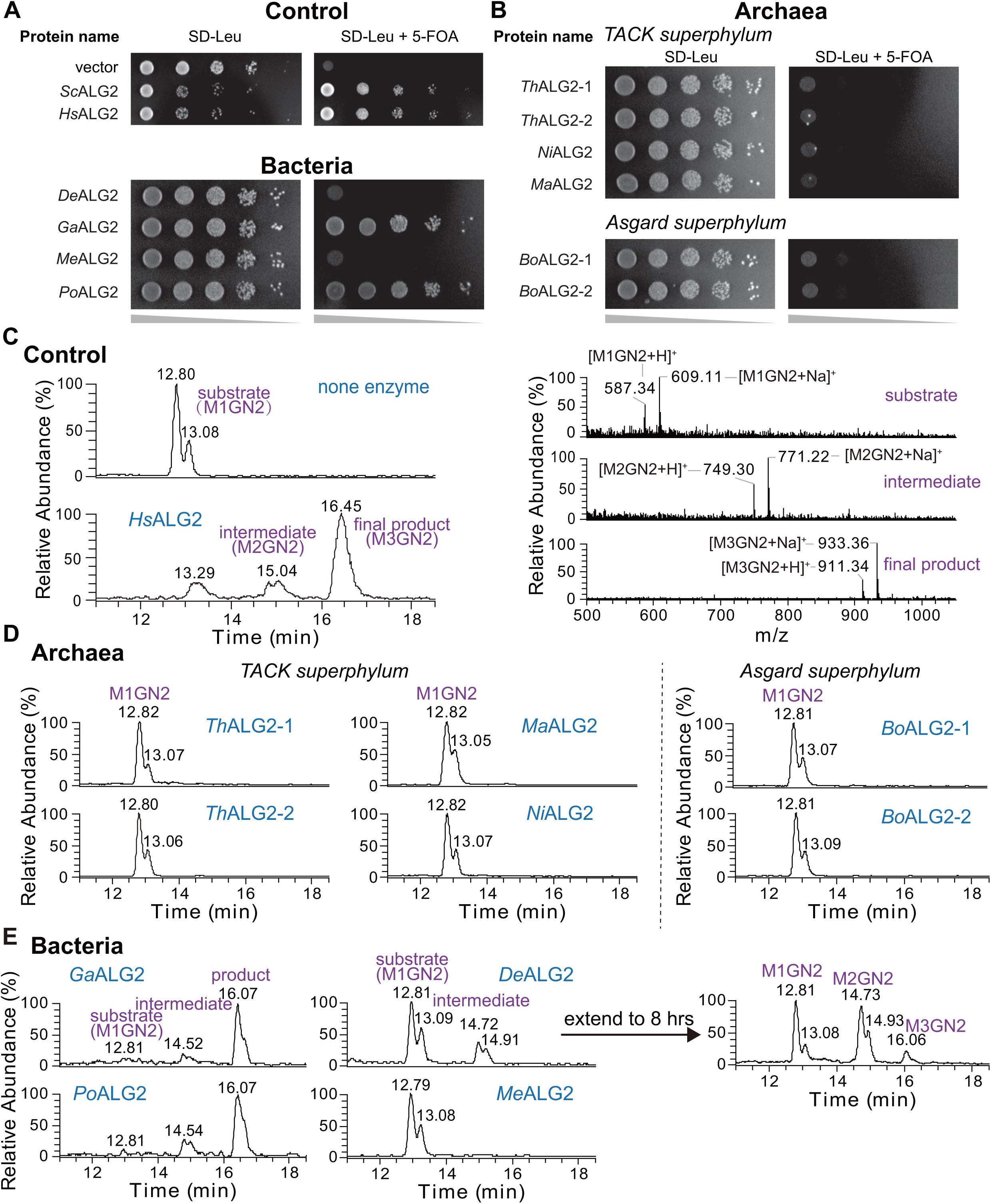
ALG2 homologs from bacteria exhibit dual 1,3 and α 1,6 MTase activity. (**A**) *LEU2* plasmids encoding *ScALG2*, *HsALG2*, and bacterial homologous sequences (including *DeALG2*, *GaALG2*, *MeALG2*, *PoALG2*), or an empty vector, were introduced into W303a-*alg2Δ* mutants carrying the *URA3*-*ScALG2* plasmid. The empty *LEU2* vector served as negative control, while *ScALG2* and *HsALG2* served as positive controls; (**B**) *LEU2* plasmids containing archaeal homologous sequences (including *ThALG2-1*, *ThALG2-2*, *NiALG2*, *MaALG2*, *BoALG2-1*, *BoALG2-2*) were transformed into W303a-*alg2Δ* mutants with the *URA3*-*ScALG2* plasmid; Yeast cells were spotted onto SD-Leu plates with or without 5-FOA and incubated at 30 °C for 48 hrs; (**C**) UPLC chromatograms of glycan chains released from ALG2 reaction mixtures. Reactions were carried out in the presence or absence of the membrane fraction of recombinant *Hs*ALG2 protein and incubated at 37 °C for 4 hrs (see *Methods* for detailed experimental conditions). Major peaks corresponding to M1GN2, M2GN2, and M3GN2 are indicated (left panel). ESI-MS spectra of glycans released from the reaction mixture containing *Hs*ALG2. The peaks eluted at ∼ 13.2 min, ∼ 15.0 min, and ∼ 16.4 min were identified as M1GN2, M2GN2, and M3GN2, respectively (right panel). *Hs*ALG2 served as a positive control for this *in vitro* assay; (**D**) UPLC chromatograms of glycan chains released from reaction assays with archaeal ALG2 homologs. The archaeal ALG2 homologs used in Figure 3B were expressed in *E. coli* cells. Membrane fractions were prepared and subjected to ALG2 activity assays under the same conditions as described in part (**C**); (**E**) UPLC chromatograms of glycan chains released from ALG2 reaction mixtures containing bacterial ALG2 homologs. The bacterial ALG2 homologs used in Figure 3C were assayed for ALG2 activity under the same conditions as the archaeal homologs in part (**B**). To further evaluate the enzymatic activity of *De*ALG2, the reaction time was extended to 8 hrs.

Eukaryotic ALG2 catalyzes addition of both the α1,3- and α1,6-linked mannose onto M1GN2-PP-Dol to form the trimannosyl core M3GN2-PP-Dol (Fig 1C). To determine if *alg2Δ* suppression is due to the α1,3- and α1,6-dual mannosyltransferase activity, recombinant protein of ALG2 homologs was directly applied to an *in vitro* ALG2 MTase assay, in which Man-(β1,4)-GN2-pyrophosphate-phytanyl (M1GN2-PP-Phy) was used as the LLO acceptor instead of the natural M1GN2-PP-Dol acceptor(Flitsch *et al*, 1992; Göker & Oren, 2023; Xiang *et al*., 2022). ALG2 homologs were expressed in *Escherichia coli* cells and confirmed by the Western-Blotting assay (Fig. EV1C). Membrane fractions from *E. coli* cells producing ALG2 homologs were added to a standard reaction and incubate with M1GN2-PP-Phy in reaction mixture at 37 ℃ for 4 hrs. After stopping the reaction, glycan products including M1GN2 (substrate), M2GN2 (intermediate), and M3GN2 (product) were chemically released from -PP-Phy and analyzed by ultra-performance liquid chromatography-mass spectrometry (UPLC-MS) (see *Methods* for details). As a positive control, *Hs*ALG2 containing membrane fraction produced three elutes around 13.2 min, 15.0 min and 16.4 min in UPLC (indicated as substrate, intermediate and final product in Fig. 3C, left panel), which possessed MS peaks at m/z: 587.34 / 609.11, 749.30 / 771.20 and 911.34 / 933.36, corresponding to [M1GN2 +H]^+^ / [M1GN2 +Na]^+^, [M2GN2 +H]^+^ / [M2GN2 +Na]^+^ and [M3GN2 +H]^+^ / [M3GN2 +Na]^+^, respectively (Fig. 3C, right panel). Noteworthy, dual peaks observed in UPLC-MS are likely the result of α- and β-anomeric forms of released glycans.

In contrast to the *Hs*ALG2, all archaeal recombinant proteins failed to convert M1GN2-PP-Phy to M2GN2-PP-Phy and M3GN2-PP-Phy science non products can be observed from their reaction mixtures (Fig. 3D). On the other hand, bacterial *Ga*ALG2 and *Po*ALG2 generated two product peaks around 14.5 and 16.0 min (Fig. 3E) in UPLC, which then been confirmed as the M2GN2 intermediate and M3GN2 final product, respectively, by MS analysis (Fig. EV1D). These results are consistent with their yeast complementation test (Fig. 3A). However, a weak dual peak eluting at the intermediate (M2GN2) position was observed from the reaction mixture with *De*ALG2 (Fig. 3E), which has failed to recover the growth defect of *alg2Δ* yeast cells (Fig. 3A). To confirm whether *De*ALG2 possesses MTase activity, we extended the incubation time from 4 to 8 hrs and analyzed the reaction mixture again. As a result, the intermediate peak was enhanced, and a new dual peak was observed at the final product (M3GN2) position (Fig. 3E). MS analysis confirmed these two peaks as the M2GN2 and M3GN2 product (Fig. EV1D), respectively, indicating the addition of mannose residues to M1GN2-PP-Phy by *De*ALG2. The failure to complement the growth defect of *alg2Δ* strain was due to its low level of activity. Taken together, our *in vitro* and *in vivo* assays confirmed functional ALG2 homologs in bacteria.

### Signature motifs required for ALG2 MTase activity are highly conserved in bacterial homologs

Previous studies have identified a highly conserved C-terminal EX_7_E and an N-terminal G-rich loop motifs in eukaryotic ALG2 protein. The EX_7_E motif works as a nucleophile that stabilizes the donor substrate (GDP-Mannose)(Kostova *et al*, 2003), while the N-terminal G-rich loop is hypothesized to function as a flexible loop required for the conformational transitions during catalysis(Steinberg, 2018). Both motifs are crucial for the MTase activity of *Sc*ALG2(Li *et al*., 2018). To seek further evidence supporting the functional homologs of ALG2 in bacteria, we revealed the evolutionary history of these two motifs.

By inspecting our MSA analysis results (Table EV4), we confirmed the N-terminal G-rich loop in all selected eukaryotic ALG2 proteins with a conserved “G-I-G-G-A-E-R” amino acid sequence (Fig. 4A). This motif can be identified in all bacterial homologs but is absent in archaea (Fig. 4A, pink box). To verify its biological importance for bacterial ALG2 functional homolog, two N-terminal glycine residues of bacterial *Ga*ALG2, including G20 and G22, were changed to alanine (A), respectively (Fig. 4B, right panel, *Ga*ALG2 mutants). As a control, similar substitutions were also introduced into *Hs*ALG2, resulting in *Hs*ALG2^G25A^ and *Hs*ALG2^G27A^ mutant proteins (Fig. 4B, left panel, *Hs*ALG2 mutants). Each of these ALG2 mutant proteins were expressed at similar levels in *E. coli* (Fig. EV2), and their membrane fractions were tested for MTase activity *in vitro*. This experiment demonstrated that both *Ga*ALG2^G20A^ and *Hs*ALG2^G25A^ completely lost their MTase activity, whereas the *Ga*ALG2^G22A^ and *Hs*ALG2^G27A^ mutations led to a significant reduction in activity (Fig. 4B). These findings indicate that the conserved N-terminal G-rich loop plays a critical role in maintaining MTase activity for bacterial *Ga*ALG2. On the other hand, the EX_7_E motif are also confirmed in all selected eukaryotic ALG2 proteins with a conserved “E-H-F-G-I-V-P-L-E” amino acid sequences (Fig. 4A). This motif can be found in all bacterial and TACK homo logs but is also absent in Asgard (Fig. 4A, cyan box). Noteworthy, in 8 of 9 archaeal (TACK) ALG2 homologs, the hydrophobic Valine (V) residue of the consensus sequence was replaced by Alanine (A) or Glycine (G) (Fig. 4A, purple dashed box). To test whether this substitution affects the MTase activity, the A residue was introduced into *Ga*ALG2 and *Hs*ALG2, resulting in the *Ga*ALG2^V317A^ and *Hs*ALG2^V330A^ mutants. Our *in vitro* assay revealed a reduction in the MTase activity of these mutant proteins compared to *Hs*ALG2 at a similar expression level (Fig. 4B, Fig EV2), suggesting the importance of this V residue for ALG2 activity.

**Figure 4.**
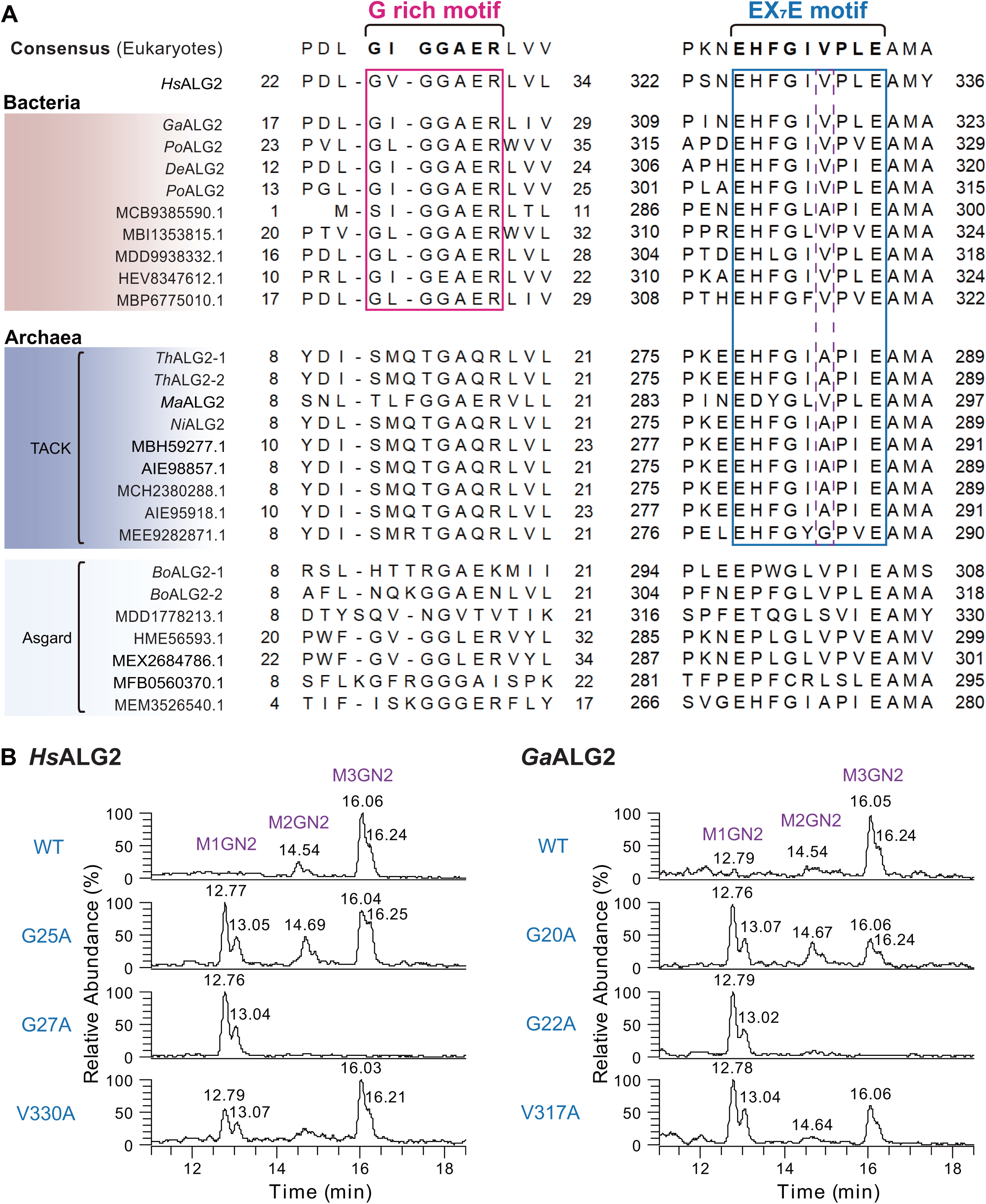
Signature motifs required for ALG2 MTase activity are conserved among bacterial homologs. (A) The multi-sequence alignment (MSA) result revealed that eukaryotic ALG2 homologs possess two characteristic motifs: a G-rich motif (G-I-G-G-A-E-R) and an EX_7_E motif (E-H-F-G-I-V-P-L-E). The positions of the amino acid residues marking the start and end of these motifs are indicated adjacent to the sequences. *Hs*ALG2 was selected as a representative sequence for eukaryotic homologs. The G-rich motif is conserved among bacterial homologs (pink box) but is absent in archaeal homologs. The EX_7_E motif is conserved in bacterial and TACK archaeal homologs (cyan box) but not in Asgard archaeal homologs. Notably, in most TACK homologs, the valine residue within this motif is substituted by alanine or glycine (purple dashed box); (**B**) The UPLC chromatograms of glycan chains released from the reaction mixtures containing either wild-type or mutant forms of *Hs*ALG2 and *Ga*ALG2 are shown. The mutants analyzed include *Hs*ALG2^G25A/G27A/V330A^ and *Ga*ALG2^G20A/G22A/V317A^. Reactions were conducted using recombinant ALG2 homolog proteins under the same conditions as described in Figure 3. Major peaks corresponding to M1GN2, M2GN2, and M3GN2 are indicated.

Taken together, these findings based on protein structure data underscore the importance of the N-terminal G-loop and the V residue of EX_7_E motif in maintaining the core function of ALG2 MTase, and well explained why bacterial, but not archaeal, ALG2 sequential homologs possess similar MTase activity with *Sc*ALG2 and *Hs*ALG2.

### Bacterial ALG1 homologs possess β −1,4 mannosyltransferase activity

As another member of cytosolic MTase complex, eukaryotic ALG1 catalyzes the addition of a β1,4-linked mannose onto GN2-PP-Dol. Our BLASTp analysis only identified reliable ALG1 homologs in bacteria but were absent in archaea (Table 1). To verify whether those bacterial ALG1 homologs have the same MTase activity, their biological activities were tested using both yeast complementation and *in vitro* ALG1 MTase assay.

Similar with the ALG2 complementary test (Fig. EV1A), bacterial *ALG1* homologous genes derived from Gammaproteobacteria (*Ga*ALG1), Polyangia (*Po*ALG1), Binatia (*Bi*ALG1), Deltaproteobacteria (*De*ALG1) and Methylomirabilota (*Me*ALG1) were cloned into a *2μ*/*LEU2* yeast expression vector and then transformed into a yeast *alg1Δ* strain, whose viability relies on the presence of a *URA3* plasmid-borne copy of *ScALG1* gene. After confirming the protein expression by Western-Blotting using anti-His tag antibody (Fig. 5A, right panel), transformants were grown on the SD-Leu plate containing 5-FOA. It was found that transformants containing *Ga*ALG1, *Po*ALG1 and *Bi*ALG1 homologs show comparable (even better) growth phenotype with that of *Hs*ALG1 (positive control), suggesting the complementation of the loss of *Sc*ALG1 (Fig. 5A, left panel).

**Figure 5.**
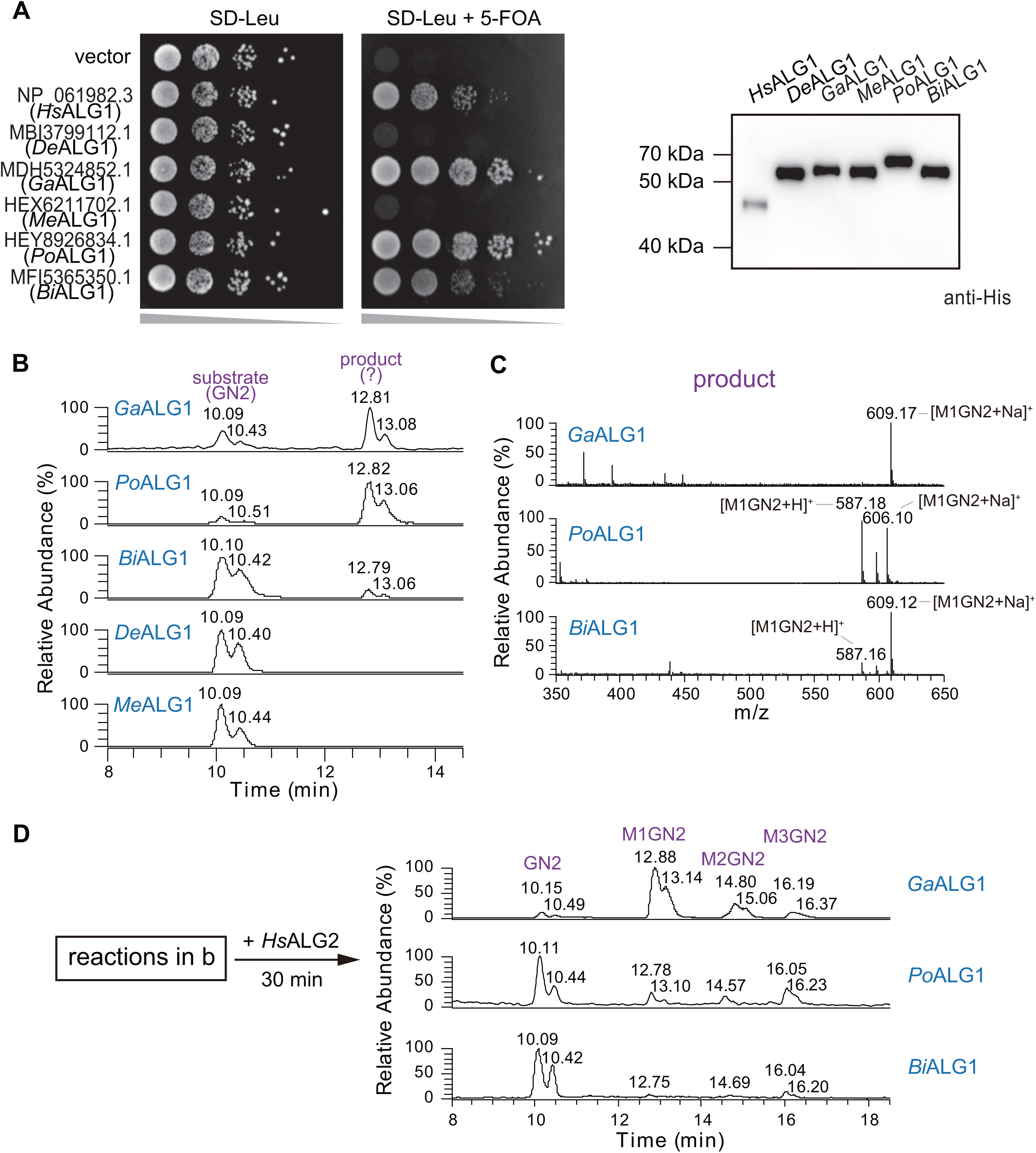
ALG1 homologs from bacteria show identical MTase activity with their yeast counterpart. (A) The principle of ALG1 yeast complementation assay is consistent with that of ALG2 in Figure 3. Subsequently, *LEU2*-based yeast expression plasmids carrying homologous *ALG1* genes from eukaryotic (*HsALG1*) and bacterial sources (including *DeALG1*, *GaALG1*, *MeALG1*, *PoALG1*, and *BiALG1*), as well as an empty vector control, were individually transformed into this yeast strain. After confirming the expression of ALG1 homologs by Western-Blotting assay, yeast viability was assessed to determine the functional complementation capacity of the homologs; (**B**) The UPLC chromatograms of glycan chains released from the reaction mixtures containing bacterial ALG1 homologs. The bacterial ALG1 homologs used in (**A**) were expressed in *E. coli* cells. Membrane fractions were prepared and incubated with the reaction buffer at 37 °C for 1.5 hrs followed by the UPLC analysis. Major peaks corresponding to substrate (GN2) and product (M1GN2?) are indicated; **c**) The ESI-MS spectra of glycans released from the reaction mixtures described in (**B**) are presented. Mass analysis reveals that the peaks eluted at ∼ 12.8 min correspond to the product (M1GN2); (**D**) The UPLC chromatogram of glycans released following the addition of *Hs*ALG2 to the reaction mixtures from (**B**), with further incubated for 30 mins, were shown.

Moreover, the above ALG2 homologs were expressed in *E. coli* cells. After confirming the expression of proteins by the Western-Blotting assay (Fig. EV3), membrane fractions producing recombinant ALG1 homologs were added to a standard reaction and incubate with GN2-PP-Phy in reaction mixture at 37 ℃. After incubating for 1.5 hrs, the reaction mixture was then divided into two halves. One of the halves was heated to stop the reaction. The glycans in the mixture including the acceptor (GN2) and products were then chemically released from -PP-Phy and analyzed by UPLC-MS. As shown in Figure 5B, the recombinant *Ga*ALG1, *Po*ALG1 and *Bi*ALG1 proteins generated a dual product peak around 12.8 and 13.1 min in UPLC, corresponding to [M1GN2 +H]^+^ / [M1GN2 +Na]^+^ (Fig. 5C), while the *De*ALG1 and *Me*ALG1 failed to do so. To confirm the linkage of mannose added by bacterial ALG1 homologs in M1GN2 products, another half of reaction were added *Hs*ALG2 and further incubated for 30 mins. It was found that M1GN2 products in ALG1 reactions were converted to M2GN2- and M3GN2- by the human ALG2, indicating these ALG1 products (M1GN2-) are indeed the substrate for human ALG2, which contain the β1,4-linked mannose (Fig. 5D). Our study confirms the existence of structural and functional ALG1 homologs in bacteria, providing another line of evidence to support the bacterial origins for cytosolic MTases.

## Discussion

N-glycosylation is a crucial post-translational modification process in eukaryotes. Investigating the evolutionary trajectories and conservation patterns of ALG GTases contributes to a deeper understanding of the evolutionary history of the N-glycosylation pathway and offers potential insights into the study of eukaryogenesis. Our study focused on the phylogenies of cytosolic ALG1 and ALG2 MTases, which are responsible for assembling the core trimannosyl M3GN2 structure—a conserved feature present in all eukaryotic N-glycans (Fig. 1C). In conjunction with functional and structural analyses, our expanded phylogenetic investigation has yielded compelling evide nce suggesting that these cytosolic ALG MTases originated from bacterial ancestors.

Phylogenies represent statistically supported hypotheses derived from available molecular, morphological, or behavioral data. Modern phylogenetic analyses primarily rely o n DNA and protein sequence databases. Despite the recent expansion of genomic data and advances in analytical methodologies, biological validation remains essential to confirm the reliability of inferred phylogenetic relationships. Furthermore, the high diversity, strict substrate specificity, and low degree of sequence conservation among ALG GTases present substantial challenges for comprehensive evolutionary comparisons across the three domains of life, thereby reinforcing the necessity of experimental validation in our study. Previous phylogenetic investigations of the N-glycosylation pathway have indicated a proteoarchaeal origin for the ALG2 MTase(Lombard, 2016). By contrast, our extended phylogenetic analysis uncovered a closer evolutionary relationship between bacterial and eukaryotic ALG2 homologs (Fig. 2). To enhance the accuracy of inference, we employed both *in vivo* and *in vitro* assays to directly evaluate the MTase activities of ALG2 orthologs. Consistent with the Bayesian clustering results, this experimental approach clearly identified dual α1,3- and α1,6-mannosyltransferase activities in bacterial ALG2 homologs, which were absent in archaeal homologs (Fig. 3A and 3B). This enzymatic profile was further corroborated by molecular analyses focusing on a conserved N-terminal G-rich loop and a C-terminal EX_7_E motif, both of which are crucial for ALG2 enzymatic function. The G-loop and a conserved valine residue within the EX_7_E motif are highly preserved in eukaryotic and bacterial homologs but not in archaeal counterparts (Fig. 4), offering structural insights into why bacterial, but not archaeal, orthologs exhibit MTase activity comparable to that of eukaryotic ALG2. Our findings present a compelling case study for comparative genomics and evolutionary biology, illustrating how the integration of phylogenetic analysis, functional assays, and structural modeling can enhance the precision and reliability of phylogenomic interpretations.

As outlined above, the closer evolutionary relationship observed between bacterial and eukaryotic ALG1 and ALG2 homologs (Fig. 2), together with the identification of their functional homologs in bacteria (Figs. 3–5), strongly supports the hypothesis that these cytosolic MTases emerged from bacterial ancestors rather than archaeal lineages. This bacterial progenitor hypothesis contradicts the conventional view that eukaryotic proteins predominantly evolved from archaeal ancestors. Historically, the prevailing hypothesis has proposed an archaeal-eukaryotic origin for the evolution of the LLO synthesis pathway(Lombard, 2016; Meyer *et al*., 2022; Raval *et al*., 2022), as archaea display more eukaryotic-like characteristics than bacteria, including dolichol-like lipids and structurally similar LLOs(Jarrell *et al*., 2014) (Fig. 1A and 1B). Furthermore, biological studies on the initiation GTase complex (Fig 1C) have provided evidence supporting a vertical inheritance model, suggesting archaeal origins for ALG7 and ALG13/14 proteins(Meyer *et al*., 2022). In contrast, this study, which focuses on ALG1 and ALG2 MTases—two subunits of the second GTase complex involved in the ER N-glycosylation pathway—reveals a distinct evolutionary pattern compared to the initiation complex, lending support to the bacterial origins for these cytosolic ALG MTases. Our findings highlight the significant contribution of bacteria to the early evolution of N-glycosylation and, most notably, provide direct evidence that the eukaryotic N-glycosylation pathway has multiple evolutionary origins.

N-glycosylation is a universally conserved post-translational modification across eukaryotes, ranging from unicellular protists to multicellular animals and plants (Aebi, 2013; Toustou *et al*., 2022). Consistent with this observation, our BLASTp analysis has revealed ALG1 and ALG2 homologs in a wide range of eukaryotic species (Table 1; Tables EV1 and EV2), indicating that they are ancient in eukaryotes. Therefore, the presence of functional ALG1 and ALG2 homologs in bacterial lineages suggests that these genes have been acquired and subsequently adapted by eukaryotic genomes. However, the precise stage at which these bacterial MTase genes integrated with those archaeal GTase genes involved in the ALG initiation complex remains unclear. Our phylogenetic analysis did not detect any ALG1 or reliable ALG2 homologs within the Asgard superphylum of archaea (Table 1; Figs. 2–5). It is conceivable to think that this integration should happen after the Asgard archaea in the evolutionary process. We propose a plausible mechanism for the genetic integration of *ALG2*, *ALG7*, and *ALG13*/*14* genes. The ancestral forms of these genes were likely present in the LUCA, after which their descendants diverged into two primary lineages: bacteria and archaea. During subsequent evolution, ALG7 and ALG13/14 homologs acquired their GTase functions within the archaeal lineage, whereas the ALG2 homolog evolved its MTase activity in bacteria, where the ALG1 MTase also emerged. These functional lineages evolved independently until their eventual convergence into a single cellular context during eukaryogenesis. We hypothesize that this integration may have occurred via an ancient horizontal gene transfer (HGT) or endosymbiotic event from bacterial donors to an archaeal host, likely at the prokaryote-eukaryote transition phase or shortly after the emergence of the Last Eukaryotic Common Ancestor (LECA). This hypothesis closely aligns with the “endosymbiotic theory”, which posits that eukaryotic cells arose through the integration of bacterial endosymbionts into archaeal hosts(Sagan, 1967). Further investigations are necessary to elucidate the underlying mechanisms and selective pressures that facilitated the integration and acquisition of *ALG1* and *ALG2* genes during eukaryogenesis. Nonetheless, our study provides a valuable framework for future studies on the evolutionary ancestry of the eukaryotic N-glycosylation pathway, with potential implications for understanding the evolution of the ER and even the process of eukaryogenesis.

## Methods

### Reagents and tools table

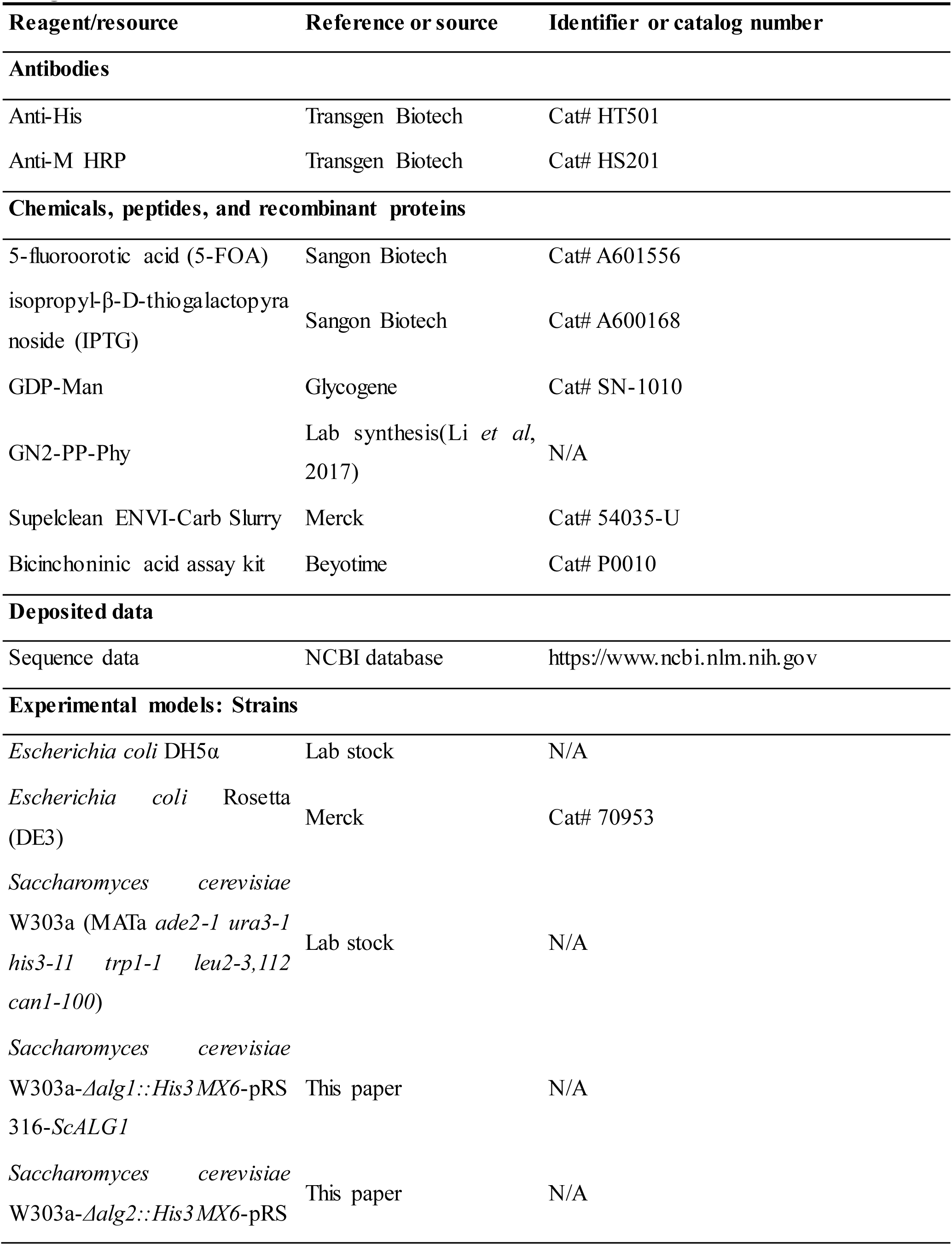
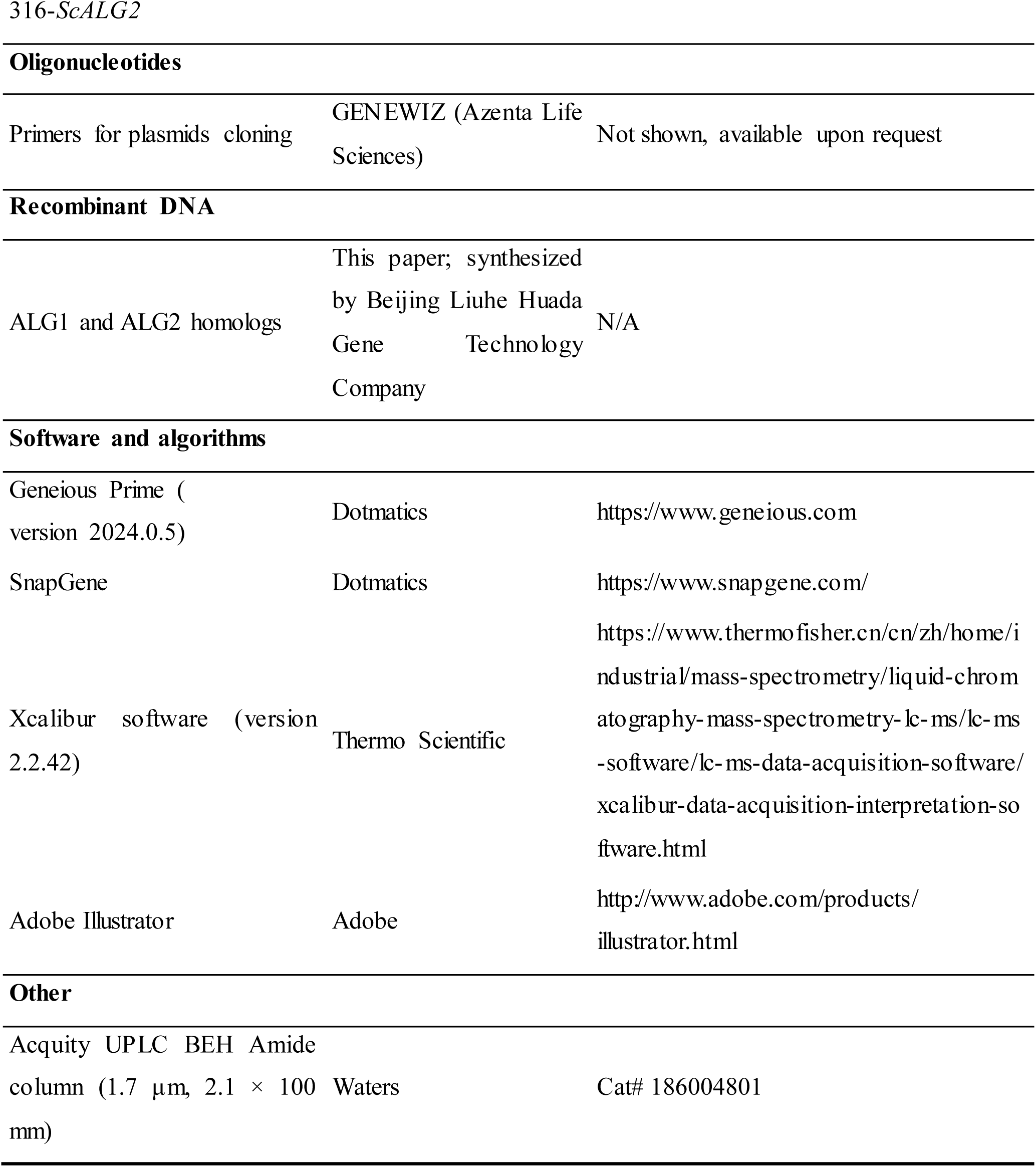

### Sequence identification and phylogenetic analysis

To identify homologs of *Hs*ALG1 and *Hs*ALG2 in prokaryotes and eukaryotes, BLASTp (https://blast.ncbi.nlm.nih.gov/Blast.cgi) were performed with the standard parameters. The E-value was cut-off at 1.00E-20 in eukaryotes, and 1.00E-7 in prokaryotes. Sequences were further treated by removing the sequences with abnormal lengths (sequences ranging from 301 to 700 amino acids in length were retained) and duplicate organism sources. To search archaeal homologs, the organism was set as “Archaea” directly. For bacterial homologs, organisms were set as 45 phyla according to the ICSP respectively and gathered(Göker & Oren, 2023; Oren & Garrity, 2021; Oren *et al*., 2022). Eukaryotic homologs were searched across four major kingdoms: Protista (including Stramenopiles, Euglenozoa, Rhizaria, Rhodophyta, Parabasalia, Heterolobosea, Dinophyceae, Cryptophyta, Perkinsea, Apicomplexa, Amoebozoa, and Ciliates), Fungi (Fungi), Plantae (Viridiplantae), and Animalia (Animalia).

For ALG2, the identified homologs were subjected to phylogenetic analysis. WbaZ from *Sinorhizobium meliloti* was used as an outgroup, as it belongs to the GT4 superfamily like ALG2 and is involved in capsular polysaccharide biosynthesis. ALG2 homolog sequences were aligned using Clustal Omega with default parameters and visualized using Geneious Prime (version 2024.0.5, https://www.geneious.com). Bayesian phylogenetic trees were constructed using the MrBayes (version 3.2.6) plugin in Geneious Prime under a Gamma substitution model.

### Plasmids, strains, and media

#### Plasmids

For expression in *E. coli*, homologous sequences were cloned into the pET32a-*TrxA*-*6His* vector. For yeast expression, *ALG1* and *ALG2* homologous sequences were inserted into the *2μ*/*LEU2* yeast shuttle vector YEp351GAPII-*6His*. All homologs can be accessed via the NCBI database (https://www.ncbi.nlm.nih.gov) by GenBank accession numbers. Standard molecular biology techniques were employed for plasmid construction. Site-directed mutagenesis was performed using overlapping PCR with mutagenic primers (primer sequences are available upon request). All plasmids were verified by DNA sequencing.

#### Strains and media

*E. coli* strain Rosetta (DE3, Merck, NJ, United States) was used for recombinant protein expression and cultured in Terrific Broth medium (1.2% tryptone, 2.4% yeast extract, 0.5% glycerol, 1.254% K_2_HPO_4_, and 0.231% KH_2_PO_4_). *Saccharomyces cerevisiae* strain W303a (MATa *ade2-1 ura3-1 his3-11 trp1-1 leu2-3,112 can1-100*) served as the parental strain. Derivative strains W303a-*alg1Δ::His3MX6*-pRS316-*ScALG1* and W303a-*alg2Δ::His3MX6*-pRS316-*ScALG2* were used for yeast complementation assays. Yeast cells were cultured in rich medium YPA (1% yeast extract, 2% peptone, 50 mg/L adenine) supplemented with 2% glucose (YPAD), or in synthetic minimal medium containing 0.67% yeast nitrogen base, and 2% glucose (SD), supplemented with appropriate amino acids.

### Yeast membrane proteins extraction and Western-Blotting analysis

For yeast membrane protein extraction, cells were lysed as previously described(Gao *et al*, 2005; Lu *et al*., 2012), and the cell lysates were centrifuged at 3,000 × *g* for 20 mins to remove unlysed cells and cell wall debris. The membrane fractions were obtained by further centrifugation (15,000 rpm, 30 mins, 4 °C), giving the pellet containing ER membrane proteins.

The expression of the proteins mentioned earlier was detected by Western-Blotting. Proteins were separated by 8% SDS-PAGE, followed by transfer onto PVDF (Bio-Rad, Hercules, CA, USA) membrane. The membrane was incubated with anti-His mouse antibody (TransGen Biotech, Beijing, China), washed, and incubated with anti-mouse IgG–horseradish peroxidase (TransGen Biotech). Immunoreactive bands were visualized by ECL (Bio-Rad).

### Yeast complementation assay

The yeast strain W303a-*alg1Δ::His3MX6*-pRS316-*ScALG1* and W303a-*alg2Δ::His3*MX6-pRS316-*ScALG2* was used to evaluate ALG1 and ALG2 homologs functionality under 5-FOA selection. *LEU2*-marked YEp351GAPII-*6His*-*ALG1* and *ALG2* homologous sequences were transformed into the respective yeast mutants and selected on SD-Leu plates at 30 °C for 48 hrs. Transformants were subsequently cultured on SD-Leu plates with or without 5-FOA at 30 °C for 48 hrs.

### Extraction of recombinant proteins expressed in *E. coli*

Recombinant *Sc*ALG1 was expressed and purified as previous study(Li *et al*., 2017). For the expression of ALG1 and ALG2 homologs, a series of *E. coli* pET32a expression plasmids were transformed into *E. coli* Rosetta (DE3) cells (Merck) and cultured for recombinant proteins. After overnight cultured of cells in 5 mL LB (Luria-Bertani) medium, 2 mL was transferred to 200 mL of TB (Terrific-Broth) medium at 37°C to an OD600 of 0.8 and then cooled to 16 °C. After adding 0.1 mM isopropyl-β-D-thiogalactopyranoside (IPTG, Sangon Biotech, Shanghai, China), the cells were incubated for additional 18 hrs at 16 °C to induce protein expression. The harvested cells were resuspended in 10 mL buffer A (150 mM NaCl, 50 mM Tris-HCl (pH 8.0)) and homogenized by sonication. The cell lysate was centrifuged at 4,000 × *g* for 20 mins to remove debris. The supernatant was further centrifuged at 15,000 rpm for 60 mins, the precipitates, which contained recombinant enzymes were resuspended in 500 μL buffer A with 1% Triton-X 100. This suspension was inverted and incubated on ice with gentle mixing for 60 mins, followed by centrifugation at 12,000 × *g* for 30 mins. The protein concentration of the supernatant was determined using a bicinchoninic acid assay kit (Beyotime, Shanghai, China). Subsequently, 30% glycerol was added, and the sample was stored at −80 °C, which was used as the crude enzyme for Western-Blotting and activity assay.

Our previous study demonstrated that the *in vitro* activity of some ALG2 homologs depend on the inclusion of membrane(Li *et al*., 2018; Xiang *et al*., 2022), so the plain bacterial membrane fraction was prepared. The empty vector pET32a was transformed into *E. coli* Rosetta (DE3) cells (Merck) harboring the empty vector pET32a. Overnight culture in 10 mL LB medium was disrupted by sonication in 500 μL buffer A, followed by centrifugation (3,000 × *g*, 20 mins) to remove cellular debris. Membranes were pelleted from the lysate by centrifugation at 15,000 rpm for 60 mins. After homogenization in 50 μ L buffer A with 30% glycerol, membrane was heated at 100 °C for 5 mins to inactivate endogenous proteins. All centrifugations were performed at 4 °C.

### In vitro *quantitative activity* assay

Recombinant ALG1 and ALG2 homologs expressed in *E. coli* were subjected to *in vitro* activity assays. The ALG1 enzyme reactions were carried out as previously described(Li *et al*., 2017) at either 30 °C or 37 °C for 1.5 hrs. The standard reaction mixture for ALG2 homologs was prepared in a final volume of 50 μ L containing: 10 mM MgCl_2_, 100 μM M1GN2-PPhy, 2 mM GDP-Man, 500 ng crude enzyme, 1 μL membrane fraction and 50 mM Tris-HCl (pH 8.0) to reach the final volume. Unless otherwise specified, reactions were conducted at 30 °C or 37 °C for 4 hrs. All reactions were terminated by heating at 100 °C for 5 mins. Prior to analysis, 20 mM HCl was added to acidify the oligosaccharides, followed by heating at 100 °C for 45 mins. The samples were then purified and lyophilized for further analysis.

### UPLC-MS analysis of oligosaccharides

Dried samples were reconstituted in water prior to UPLC-MS analysis. Samples were injected into a Dionex Ultimate 3000 UPLC system (Thermo Scientific, MA, USA). Glycans were separated on an Acquity UPLC BEH Amide column (1.7 μm, 2.1 × 100 mm, Waters, MA, USA) using an acetonitrile gradient at a flow rate of 0.2 mL/min. The gradient program was as follows: 0–2 mins, isocratic 80% acetonitrile; 2–15 mins, linear decrease to 50% acetonitrile; 15–20 mins, isocratic 50% acetonitrile. Eluted glycans were analyzed by electrospray ionization mass spectrometry (ESI-MS) using a TSQ Quantum Ultra (Thermo Scientific, MA, USA) in positive ion mode across a mass range of 300 – 2000 m/z. Oligosaccharide transfer efficiency was quantified by measuring peak intensities in the LC-ESI-MS data using Xcalibur software (version 2.2.42, Thermo Scientific, USA).

## Acknowledgements

We are grateful to members of the Gao laboratory for reagents, comments and other contributions to this project.

## Funding

This work was supported by grants-in-aid from the National Natural Science Foundation of China (32271342).

## Author contributions

Conceptualization, X.-D.G., and H.N.; methodology, S.X., S.C., and Y.-H.G.; validation, S.X., and Y.-F.H.; formal analysis, S.X., S.C., and Y.-H.G.; investigation, S.X., Y.-F.H., and X.-D.G.; writing—original draft preparation, S.X., S.C., and X.-D.G.; writing—review and editing, H.N., and X.-D.G.; supervision, X.-D.G., and H.N.; funding acquisition, X.-D.G., and H.N. All authors have read and agreed to the published version of the manuscript.

## Disclosure and competing interests statement

The authors declare no conflict of interest.

---

**Figure EV1. Data supporting Figure 3.**
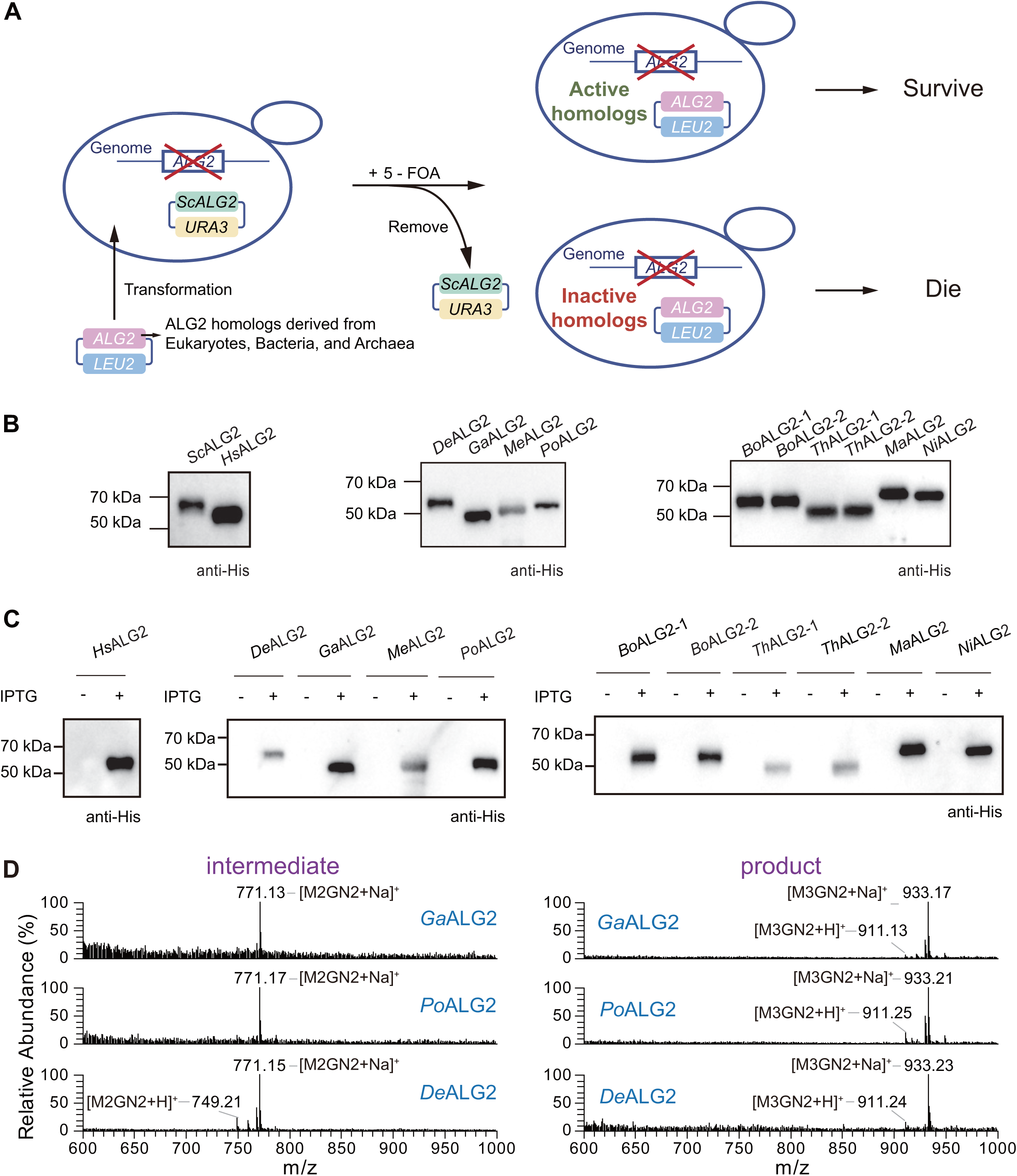
Recombinant prokaryotic ALG2 homologs expression analysis. (**A**) A schematic representation illustrates the use of 5-FOA in the yeast complementation assay. A *URA3*-*ScALG2* plasmid was introduced into the W303a yeast strain, from which the endogenous *ALG2* gene had been deleted. *LEU2*-based yeast expression plasmids harboring individual *ALG2* homologous genes were transformed into this strain. After removing the *URA3* plasmid by 5-FOA treatment, the yeast viability was then assessed to determine the functional complementation capacity of the homologs; (**B**) Expression of the ALG2 homologs using for yeast complementary assay in Figure 3A and B was confirmed by SDS-PAGE followed by immunoblotting using an anti-His monoclonal antibody; (**C**) pET32a plasmids encoding *Hs*ALG2 and prokaryotic *ALG2* homologous sequences were transformed to *E. coli* Rosetta strain, and induced expression by IPTG (see *Methods* for detailed experimental conditions). The samples were collected before and after induction, and confirmed the expression of recombinant proteins b y SDS-PAGE followed by immunoblotting using an anti-His monoclonal antibody; (**D**) ESI-MS spectra of glycans released from the reaction mixture shown in Figure 3E. The peaks eluted at ∼ 14.5 min and ∼ 16.0 min correspond to the intermediate (M2GN2) and the product (M3GN2), respectively.

**Figure EV2. Data supporting Figure 4.**
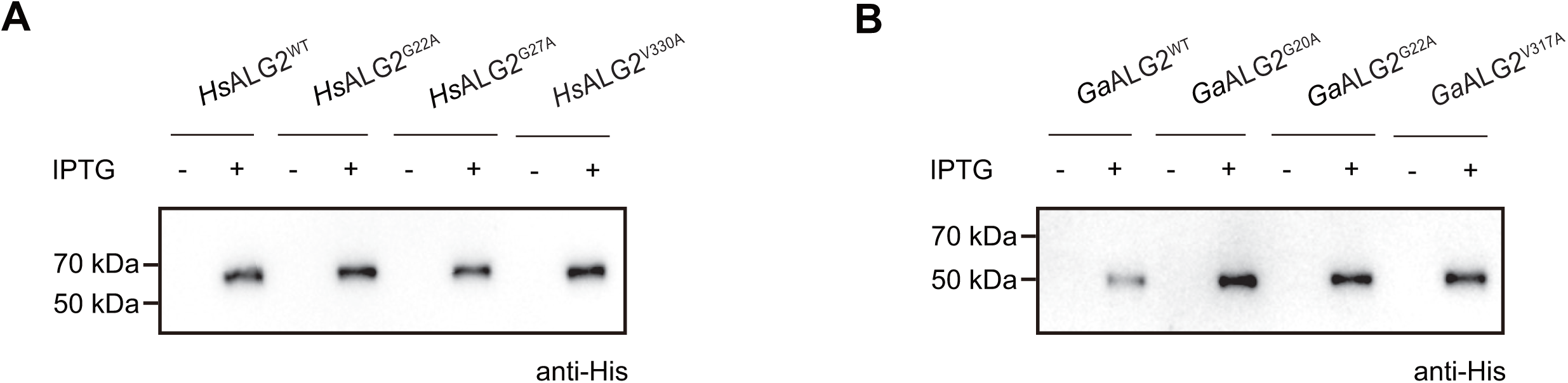
Recombinant wild-type and mutants of *Hs*ALG2 and *Ga*ALG2 homologs expression analysis. (**A**, **B**) pET32a plasmids encoding wild-type and mutants of *HsALG2* and *GaALG2* homologous sequences were transformed to *E. coli* Rosetta strain, and induced expression by IPTG (see *Methods* for detailed experimental conditions). The samples were collected before and after induction, and confirmed the expression of recombinant proteins by SDS-PAGE followed by immunoblotting using an anti-His monoclonal antibody. (**A**) Recombinant wild-type and mutants of *Hs*ALG2 expression analysis. (**B**) Recombinant wild-type and mutants of *Ga*ALG2 expression analysis.

**Figure EV3. Data supporting Fig. 5.**
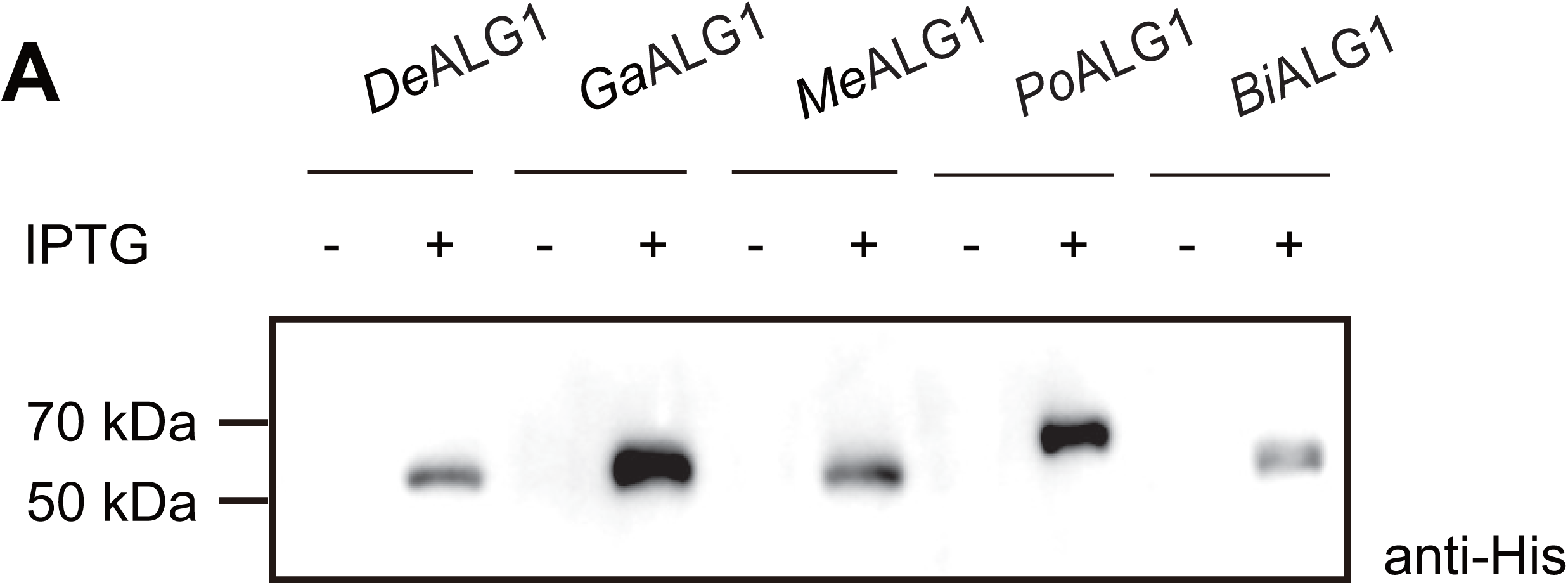
Recombinant bacterial ALG1 homologs expression analysis. **(A)** pET32a plasmids encoding prokaryotic *ALG1* homologous sequences (including *DeALG1*, *GaALG1*, *MeALG1*, *PoALG1*, and *BiALG1*) were transformed to *E. coli* Rosetta strain, and induced expression by IPTG (see *Methods* for detailed experimental conditions). The samples were collected before and after induction, and confirmed the expression of recombinant proteins by SDS-PAGE followed by immunoblotting using an anti-His monoclonal antibody.

**Table EV1. The result statistics of *Hs*ALG2 BLAST.** Different worksheets represent different kingdoms (including Bacteria, Archaea, Protista, Fungi, Plantae, and Animalia).

**Table EV2. The result statistics of *Hs*ALG1 BLAST.** Different worksheets represent different kingdoms (including Bacteria, Archaea, Protista, Fungi, Plantae, and Animalia).

**Table EV3. Sequences information of ALG2 homologs used in phylogenetic analysis**

**Table EV4. Multi-sequence alignment result of ALG2 homologs used in phylogenetic analys is**

## References

Abu-Qarn M, Yurist-Doutsch S, Giordano A, Trauner A, Morris HR, Hitchen P, Medalia O, Dell A, Eichler J (2007) *Haloferax volcanii* AglB and AglD are involved in N-glycosylation of the S-layer glycoprotein and proper assembly of the surface layer. J Mol Biol 374: 1224–1236

Aebi M (2013) N-linked protein glycosylation in the ER. Bba-Mol Cell Res 1833: 2430–2437

Aebi M, Bernasconi R, Clerc S, Molinari M (2010) N-glycan structures: recognition and processing in the ER. Trends Biochem Sci 35: 74–82

Chen S, Pei CX, Xu S, Li HJ, Liu YS, Wang YC, Jin C, Dean N, Gao XD (2024) Rft1 catalyzes lipid-linked oligosaccharide translocation across the ER membrane. Nat Commun 15

Cherepanova N, Shrimal S, Gilmore R (2016) N-linked glycosylation and homeostasis of the endoplasmic reticulum. Curr Opin Cell Biol 41: 57–65

Drula E, Garron ML, Dogan S, Lombard V, Henrissat B, Terrapon N (2022) The carbohydrate-active enzyme database: functions and literature. Nucleic Acids Res 50: D571–D577

Flitsch SL, Pinches HL, Taylor JP, Turner NJ (1992) Chemo-enzymatic synthesis of a lipid-linked core trisaccharide of *N*-linked glycoproteins. Journal of the Chemical Society, Perkin Transactions 1 : 2087–2093

Fujinami D, Taguchi Y, Kohda D (2017) Asn-linked oligosaccharide chain of a crenarchaeon, *Pyrobaculum calidifontis*, is reminiscent of the eukaryotic high-mannose-type glycan. Glycobiology 27: 701–712

Gao XD, Nishikawa A, Dean N (2004) Physical interactions between the Alg1, Alg2, and Alg11 mannosyltransferases of the endoplasmic reticulum. Glycobiology 14: 559–570

Gao XD, Tachikawa H, Sato T, Jigami Y, Dean N (2005) Alg14 recruits Alg13 to the cytoplasmic face of the endoplasmic reticulum to form a novel bipartite UDP-N-acetylglucosamine transferase required for the second step of N-linked glycosylation. J Biol Chem 280: 36254–36262

Göker M, Oren A (2023) Valid publication of four additional phylum names. Int J Syst Evol Micr 73

Helenius J, Ng DTW, Marolda CL, Walter P, Valvano MA, Aebi M (2002) Translocation of lipid-linked oligosaccharides across the ER membrane requires Rft1 protein. Nature 415: 447–450

Jarrell KF, Ding Y, Meyer BH, Albers SV, Kaminski L, Eichler J (2014) N-Linked Glycosylation in Archaea: a Structural, Functional, and Genetic Analysis. Microbiol Mol Biol R 78: 304–341

Kawakami N, Fujisaki S (2018) Undecaprenyl phosphate metabolism in Gram-negative and Gram-positive bacteria. Biosci Biotech Bioch 82: 940–946

Kifer D, Čorak N, Domazet-Lošo M, Kasalo N, Lauc G, Klobučar G, Domazet-Lošo T (2025) Contrasting macroevolutionary patterns in the human N-glycosylation pathway. Engineering

Knauer R, Lehle L (1999) The oligosaccharyltransferase complex from yeast. Bba-Gen Subjects 1426: 259–273

Kostova Z, Yan BC, Vainauskas S, Schwartz R, Menon AK, Orlean P (2003) Comparative importance *in vivo* of conserved glutamate residues in the EX_7_E motif retaining glycosyltransferase Gpi3p, the UDP-GlcNAc-binding subunit of the first enzyme in glycosylphosphatidylinositol assembly. Eur J Biochem 270: 4507–4514

Kowarik M, Numao S, Feldman MF, Schulz BL, Callewaert N, Kiermaier E, Catrein I, Aebi M (2006) N-linked glycosylation of folded proteins by the bacterial oligosaccharyltransferase. Science 314: 1148–1150

Li ST, Wang N, Xu S, Yin J, Nakanishi H, Dean N,Gao XD (2017) Quantitative study of yeast Alg1 beta-1, 4 mannosyltransferase activity, a key enzyme involved in protein N-glycosylation. Biochim Biophys Acta Gen Subj 1861: 2934–2941

Li ST, Wang N, Xu XX, Fujita M, Nakanishi H, Kitajima T, Dean N, Gao XD (2018) Alternative routes for synthesis of N-linked glycans by Alg2 mannosyltransferase. The FASEB Journal 32: 2492–2506

Liu Y, Makarova KS, Huang WC, Wolf YI, Nikolskaya AN, Zhang XX, Cai MW, Zhang CJ, Xu W, Luo ZH et al (2021) Expanded diversity of Asgard archaea and their relationships with eukaryotes. Nature 593: 553–557

Lombard J (2016) The multiple evolutionary origins of the eukaryotic *N*-glycosylation pathway. Biol Direct 11

Lu J, Takahashi T, Ohoka A, Nakajima K-i, Hashimoto R, Miura N, Tachikawa H, Gao X-D (2012) Alg14 organizes the formation of a multiglycosyltransferase complex involved in initiation of lipid-linked oligosaccharide biosynthesis. Glycobiology 22: 504–516

Meyer BH, Adam PS, Wagstaff B, Kolyfetis GE, Probst AJ, Albers SV, Dorfmueller HC, Cole PA (2022) Agl24 is an ancient archaeal homolog of the eukaryotic N-glycan chitobiose synthesis enzymes. Elife 11

Meyer BH, Shams-Eldin H, Albers SV (2017) AglH, a thermophilic UDP-*N*-acetylglucosamine-1-phosphate: dolichyl phosphate GlcNAc-1-phosphotransferase initiating protein *N*-glycosylation pathway in *Sulfolobus acidocaldarius*, is capable of complementing the eukaryal Alg7. Extremophiles 21: 121–134

Mikusova K, Mikus M, Besra GS, Hancock I, Brennan PJ (1996) Biosynthesis of the linkage region of the mycobacterial cell wall. Journal of Biological Chemistry 271: 7820–7828

Mitachi K, Siricilla S, Yang D, Kong Y, Skorupinska-Tudek K, Swiezewska E, Franzblau SG, Kurosu M (2016) Fluorescence-based assay for polyprenyl phosphate-GlcNAc-1-phosphate transferase (WecA) and identification of novel antimycobacterial WecA inhibitors. Anal Biochem 512: 78–90

Mohorko E, Glockshuber R, Aebi M (2011) Oligosaccharyltransferase: the central enzyme of N-linked protein glycosylation. J Inherit Metab Dis 34: 869–878

Noffz C, Keppler-Ross S, Dean N (2009) Hetero-oligomeric interactions between early glycosyltransferases of the dolichol cycle. Glycobiology 19: 472–478

Nothaft H, Szymanski CM (2010) Protein glycosylation in bacteria: sweeter than ever. Nat Rev Microbiol 8: 765–778

O’Reilly MK, Zhang GF, Imperiali B (2006) In vitro evidence for the dual function of Alg2 and Alg11: Essential mannosyltransferases in N-linked glycoprotein biosynthesis. Biochemistry-Us 45: 9593–9603

Oren A, Garrity GM (2021) Valid publication of the names of forty-two phyla of prokaryotes. Int J Syst Evol Micr 71

Oren A, Mares J, Rippka R (2022) Validation of the names *Cyanobacterium* and *Cyanobacterium stanieri*, and proposal of *Cyanobacteriota* phyl. nov. Int J Syst Evol Micr 72

Raval PK, Garg SG, Gould SB (2022) Endosymbiotic selective pressure at the origin of eukaryotic cell biology. Elife 11

Sagan L (1967) On the origin of mitosing cells. J Theor Biol 14: 255–274

Schmaltz RM, Hanson SR, Wong CH (2011) Enzymes in the Synthesis of Glycoconjugates. Chem Rev 111: 4259–4307

Schwarz F, Aebi M (2011) Mechanisms and principles of N-linked protein glycosylation. Curr Opin Struc Biol 21: 576–582

Shams-Eldin H, Chaban B, Niehus S, Schwarz RT, Jarrell KF (2008) Identification of the archaeal *alg7* gene homolog (encoding *N*-acetylglucosamine-1-phosphate transferase) of the N-linked glycosylation system by cross-domain complementation in *Saccharomyces cerevisiae*. J Bacteriol 190: 2217–2220

Shrimal S, Gilmore R (2019) Oligosaccharyltransferase structures provide novel insight into the mechanism of asparagine-linked glycosylation in prokaryotic and eukaryotic cells. Glycobiology 29: 288–297

Spang A, Saw JH, Jorgensen SL, Zaremba-Niedzwiedzka K, Martijn J, Lind AE, van Eijk R, Schleper C, Guy L, Ettema TJG (2015) Complex archaea that bridge the gap between prokaryotes and eukaryotes. Nature 521: 173–179

Steinberg SF (2018) Post-translational modifications at the ATP-positioning G-loop that regulate protein kinase activity. Pharmacological Research 135: 181–187

Toustou C, Walet-Balieu M-L, Kiefer-Meyer M-C, Houdou M, Lerouge P, Foulquier F, Bardor M (2022) Towards understanding the extensive diversity of protein *N*-glycan structures in eukaryotes. Biological Reviews 97: 732–748

van Wolferen M, Shajahan A, Heinrich K, Brenzinger S, Black IM, Wagner A, Briegel A, Azadi P, Albers SV (2020) Species-Specific Recognition of *Sulfolobales* Mediated by UV-Inducible Pili and S-Layer Glycosylation Patterns. Mbio 11

Wacker M, Linton D, Hitchen PG, N ita-Lazar M, Haslam SM, North SJ, Panico M, Morris HR, Dell A, Wren BW et al (2002) N-linked glycosylation in *Campylobacter jejuni* and its functional transfer into *E.coli*. Science 298: 1790–1793

Weerapana E, Imperiali B (2006) Asparagine-linked protein glycosylation: from eukaryotic to prokaryotic systems. Glycobiology 16: 91–101

Xiang MH, Xu XX, Wang CD, Chen S, Xu S, Xu XY, Dean N, Wang N, Gao XD (2022) Topological and enzymatic analysis of human Alg2 mannosyltransferase reveals its role in lipid-linked oligosaccharide biosynthetic pathway. Commun Biol 5

Yurist-Doutsch S, Chaban B, VanDyke DJ, Jarrell KF, Eichler J (2008) Sweet to the extreme: Protein glycosylation in Archaea. Mol Microbiol 68: 1079–1084

Zaremba-Niedzwiedzka K, Caceres EF, Saw JH, Bäckström D, Juzokaite L, Vancaester E, Seitz KW, Anantharaman K, Starnawski P, Kjeldsen KU et al (2017) Asgard archaea illuminate the origin of eukaryotic cellular complexity. Nature 541: 353–358

